# The impact of seizures on REM sleep and the cholinergic pedunculopontine nucleus in a mouse model of Dravet Syndrome

**DOI:** 10.64898/2026.02.17.706417

**Authors:** Krystal M. Santiago-Colon, Chandni Rana, Brandon Toth, Julia A. Kravchenko, Joseph Barden, Isabella VanHorn, Alexandra Jack, Alexander Hollier, Rida Qureshi, Hudson Haberland, Juan Disla, Nagham Eldroubi, Saif Siddiqui, Sophie Erb-Watson, Christian Burgess, Lori L. Isom, Joanna Mattis

**Affiliations:** Department of Pharmacology, University of Michigan, Ann Arbor, MI USA; Neuroscience Graduate Program, University of Michigan, Ann Arbor, MI USA; Department of Neurology, University of Michigan, Ann Arbor, MI USA; Michigan Neuroscience Institute, University of Michigan, Ann Arbor, MI USA; Department of Molecular and Integrative Physiology, University of Michigan, Ann Arbor, MI USA; VA Ann Arbor Healthcare System, Ann Arbor, Michigan, USA

## Abstract

Sleep disruption is a common and burdensome feature in epilepsy, with rapid eye movement (REM) sleep particularly affected. While sleep disturbances in epilepsy patients are multifactorial, clinical evidence suggests that recent seizures acutely impair REM sleep architecture. To investigate this relationship, we used a haploinsufficient mouse model of Dravet Syndrome, which allows experimental control of seizure timing and burden. We found that hyperthermia-induced seizures profoundly decreased subsequent sleep, specifically impairing REM entry. *In vivo* fiber photometry revealed acute, seizure-induced activation of cholinergic neurons in the pedunculopontine nucleus (PPN), a brainstem structure critical for REM entry. We additionally found that repeated seizures triggered anatomical changes in the PPN, including reduced cholinergic neuron number and significant hypertrophy of remaining cholinergic neurons. These results suggest seizures are a driver of both acute and chronic disruption of PPN cholinergic networks, which in turn impair REM sleep in epilepsy. Our findings identify the PPN as a potential therapeutic target for interventions to address sleep-related sequelae of seizures.

## Introduction

The associations between sleep and epilepsy are profound and bidirectional: sleep deprivation and sleep state strongly modify seizure risk (e.g.,(Bazil and Walczak, 1997; Nakken et al., 2005)), and in turn, sleep is frequently impaired in patients with epilepsy. Rapid eye movement (REM) sleep – during which crucial synaptic pruning and memory consolidation occurs (Peever and Fuller, 2016; MacDonald and Cote, 2021) – is particularly impacted in epilepsy. For instance, among patients admitted to an Epilepsy Monitoring Unit for diagnostic purposes, the percentage of REM sleep (REM %) was strongly reduced in those who ultimately received an epilepsy diagnosis (Sadak et al., 2022).

Causes of sleep disruption in patients with epilepsy are often multifactorial, as there are high rates of obstructive sleep apnea in this patient population (Malow et al., 2000) and anti-seizure medications may impact sleep (Jain and Glauser, 2014; Ye et al., 2022). However, intriguing clinical data provide evidence that seizures themselves directly interfere with REM sleep. On an acute timescale, multiple studies correlating seizure timing with subsequent sleep architecture (in patients with known epilepsy) found impaired REM sleep specifically in those who experienced a seizure prior to the sleep period of interest (Bazil et al., 2000; Kilgore-Gomez et al., 2024; Kremen et al., 2026). On a longer timescale, patients who underwent surgery for refractory epilepsy reported improved sleep quality (Carrion et al., 2010) and exhibited significantly increased total sleep time – particularly of REM sleep – even in the absence of significant medication changes that could confound the results (Serafini et al., 2012).

Although impaired sleep subjectively worsens quality of life in patients with epilepsy (e.g., (Wu et al., 2026)) and REM sleep disruption has numerous cognitive and behavioral consequences (e.g., (Pesonen et al., 2024), there are currently no targeted treatments available to prevent or reverse the development of this sequela of epilepsy. Progress on this front is limited by a lack of understanding of the acute and chronic impact of seizures on relevant brain structures, including the pedunculopontine nucleus (PPN), a cholinergic nucleus in the upper brainstem that regulates REM sleep entry (Shiromani and McGinty, 1986; Webster and Jones, 1988; Van Dort et al., 2015).

In this study, we sought to address this gap by characterizing the impact of seizures on sleep architecture and on cholinergic PPN neurons in a preclinical mouse model of Dravet syndrome (DS) (Mistry et al., 2014). We selected this model because DS mice have a well-characterized epilepsy phenotype – with spontaneous seizures starting at P14-P18 (Hawkins et al., 2019; Han et al., 2020) – but also have seizures that can be readily induced by exposure to hyperthermia (Oakley et al., 2009; Hawkins et al., 2017), enabling experimental control over seizure burden and timing. We found that hyperthermic seizure induction in DS mice impaired subsequent REM sleep, acutely activated PPN cholinergic neurons, and triggered anatomic changes within the PPN.

## Methods

### Experimental animals

All animal use was approved by the Institutional Animal Care & Use Committee (IACUC) at the University of Michigan. Animals were housed in same-sex cages of up to 5 mice on a 12-hour light/dark cycle and allowed food and water ad libitum. Male and female mice were used in equal proportions.

The hemizygous *Scn1a* mouse model (*Scn1a^+/-^*; RRID:MMRRC_037107-JAX) contains a targeted deletion of the first exon of the *Scn1a* gene that is associated with a ∼50% decrease in Nav1.1 protein (Miller et al., 2014; Mistry et al., 2014). *Scn1a^+/-^* mice were maintained on a 129S6.SvEvTac background. All experimental animals were bred to the 50:50 129S: BL6/J background on which the DS phenotype has been extensively characterized (Miller et al., 2014; Mistry et al., 2014; Hawkins et al., 2017; Han et al., 2020). Thus, to generate F1 experimental DS mice, F0 male *Scn1a^+/-^* mice on the 129S6.SvEvTac background were bred with F0 female mice on a C57BL/6J background, as detailed below.

For experiments that did not require selective targeting or endogenous labeling of cholinergic neurons, F0 females were wildtype (WT) C57BL/6J mice (RRID:IMSR_JAX:000664). For experiments requiring selective targeting of cholinergic neurons, F0 females were knock-in *Chat^tm2(cre)Lowl/^J* (ChAT-Cre) mice (RRID:IMSR_JAX:006410), in which Cre recombinase is expressed within choline acetyltransferase (ChAT)-expressing neurons (Rossi et al., 2011). For experiments requiring endogenous labeling of cholinergic neurons, F0 females were ChAT-Cre;tdT double-transgenic mice, in which tdTomato is expressed within cholinergic neurons; these mice were generated by crossing ChAT-Cre knock-in mice with the tdTomato reporter/Ai14 strain (Madisen et al., 2010) (RosaCAG-LSL-tdTomato; RRID:IMSR_JAX:007914), both of which were bred to homozygosity such that 100% of the F1 progeny (from the subsequent *Scn1a^+/-^*cross) were heterozygous for both Cre and tdT.

All mice were genotyped using PCR analysis of toe tissue obtained via distal phalanx amputation at postnatal day (P)5-10. WT (*Scn1a^+/+^*) littermates from each cross were used as controls.

### Seizure induction

To evoke seizures in DS mice, animals were placed in a closed plexiglass chamber (LSA Scientific Instrument Shop) under a heating lamp (Physitemp). For simultaneous fiber photometry/EEG recording, mice were additionally head-fixed using a custom apparatus (ThorLabs). Experimental animals were heated until the mouse exhibited a behavioral seizure or until the mouse internal body temperature – as measured with a rectal probe (Physitemp) – reached 42.5°C. Animals were then actively cooled using ice until their internal body temperature recovered to at least 39°C. For stereology and slice electrophysiology experiments, mice were heated in pairs (when possible), such that each DS mouse was paired with a sex-matched WT littermate control, and both mice were removed from the heating chamber when the DS mouse exhibited a seizure.

To generate positive control tissue for immunostaining (see below), status epilepticus was induced in WT (129S6.SvEvTac background) mice via intraperitoneal injection of 10 mg/kg kainic acid (Hello Bio Inc, Cat# HB0355). Mice were perfused as described below, 24 hours post-injection.

### Stereotactic surgeries

Mice were anesthetized by continuous inhalation of 1-2% isoflurane in oxygen. Once deeply anesthetized (as determined by toe pinch), mice were secured to a stereotactic apparatus (Kopf Instruments). The skull was leveled by adjusting the head position until the difference in height between bregma and lambda was < 0.05 mm. Craniotomies were made using an automatic drill (Kopf Instruments).

Viruses were administered using a 10 µL syringe and a 33-gauge beveled needle (Hamilton). For expression of Cre-dependent GCaMP8s, we injected AAV9-syn-FLEX-jGCaMP8s-WPRE (Addgene). Infusions of 300 nL (2.7 x 10^13^ genetic copies/ mL) were targeted to the right PPN at two different depths (relative to Bregma: A/P: −4.8 mm; M/L: 1.25 mm; D/V: −3.65 (100 nL) and −3.5 mm (200 nL)). All volumes were infused at a rate of 100 nL/minute, and the needle was left in place for 5 minutes before withdrawal.

For fiber photometry experiments, a 400 µm diameter fiber optic (Thorlabs) was implanted over the right PPN (A/P: −4.6 mm, M/L: 1.2 mm, D/V: −3.5 mm). For electrocorticography recordings, stainless steel skull screws (P1 Technologies) were implanted over the left temporal cortex (A/P: −2.0 mm, M/L: −2.0 mm) and right frontal cortex (A/P: 1.5 mm, M/L: 1.5 mm), with a reference electrode over the cerebellum (A/P: −5.0 mm, M/L: −1.25 mm).

For electromyography recordings, 3 stainless steel wires coated in Teflon were inserted into the nuchal muscles. 1 mm of Teflon coating was stripped to expose bare wires to muscle tissue.

Mice were also implanted with a head post (H.E. Parmer Co.) to enable head-fixed recordings. All implants were secured to the skull with a layer of dental cement (C&B Metabond) followed by dental acrylic (Fastray).

### Polysomnography

Beginning 2-3 weeks after surgery, mice were connected to an EEG/EMG recording tether and a fiber optic patch cable and given 5-7 days to habituate to the recording environment. Mice were considered fully habituated when observed to have built a nest and to comfortably use the in-cage food hopper, lickspout, and running wheel. Mice were able to move freely within the behavior chamber while outfitted with both optic fibers and EEG/EMG tethers.

Polysomnographic signals were digitized at 1000 Hz, with a 0.3-100 Hz bandpass filter applied to the EEG and a 30-100 Hz bandpass filter applied to the EMG (ProAmp-8, CWE Inc.), with a National Instruments data acquisition card and collected using a custom MATLAB script. EEG/EMG signals were notch filtered at 60 Hz to attenuate line noise. Prior to analysis, EEG recordings were high-pass filtered at 1 Hz to remove low-frequency artifacts.

Polysomnographic signals were analyzed using AccuSleep, an open-source sleep scoring algorithm in MATLAB, and verified by manual inspection. Behavioral states were scored in 5-second epochs as either wake, rapid eye movement (REM), or non-REM (NREM) sleep.

Statistical analyses of the above quantifications were performed using GraphPad Prism version 10.4.1 for Windows (GraphPad Software, Boston, Massachusetts, USA; www.graphpad.com). GraphPad Prism was also used to generate all associated graphs.

### Electrocorticography (EEG) data collection and analysis for seizure experiments

EEG signals were acquired and digitized with an Open Ephys data acquisition board (DAQ) at a sampling frequency of 1000 Hz. The raw signal was band-pass filtered from 1-70 Hz and notch filtered at 60 Hz to attenuate line noise. The processed signal was used to visually determine precise peri-ictal timepoints. The unequivocal electrographic onset (UEO) was defined as the beginning of a disruption in background EEG activity, consisting of spike-wave discharges and/or an evolution in frequency or amplitude of rhythmic activity lasting at least 10 seconds in duration. The earliest electrographic change (EEC) preceded the UEO and was defined as the time of the earliest pre-ictal spike within 1 minute of the UEO. If no pre-ictal spikes were observed, the EEC was defined at the same time point as the UEO. Seizures were consistently followed by a post-ictal attenuation, which was used to determine seizure offset (OFF).

A custom detection pipeline was developed in Python using MNE-Python and SciPy to detect prolonged high-amplitude events in a single EEG channel sampled at 500 Hz, with no resampling applied. Recordings were processed with a 60 Hz notch filter and a fourth-order Butterworth bandpass filter (1-70 Hz), after which signals were standardized via z-scoring. A threshold defined by the 97th percentile of the z-values was applied for every non-overlapping one-second segment, and segments were marked as active when at least 10% of samples exceeded this threshold. Runs of active segments lasting seven seconds or longer were labeled as candidate seizures; adjacent events separated by 30 seconds or less were combined. All events were visualized for manual verification.

### Fiber photometry data collection and analysis

For recordings performed concurrently with polysomnography, LEDs (Plexon; 473 nm) were set such that a light intensity of < 0.1 mW entered the brain; light intensity was kept constant across sessions. Emission light was passed through a filter cube (Doric) before being focused onto a sensitive photodetector (2151, Newport). Recordings were 12 hours in length and contained the last half of the dark cycle (8 PM − 2 AM) and the first half of the light cycle (2 AM − 8 AM). Only transitions in which at least 20 s of a state occurred on either side of the transition were included.

For all other recordings, fiber photometry signals were acquired at 100 Hz using a fluorescence excitation/detection module (Doric) connected to a PyPhotometry acquisition board. Calcium-sensitive fluorescence from GCaMP (470 nm) and its isosbestic point (405 nm) were independently acquired to perform offline motion correction, artifact rejection, photobleaching, etc. Raw signals were up-sampled to 1000 Hz and aligned to the EEG. Both signals were then Gaussian filtered (σ = 75, kernel size = 3σ) for denoising. The motion corrected normalized fluorescence (ΔF/F) was calculated with the equation: (GCaMP signal – isosbestic signal) / isosbestic signal. Corrected traces were then z-scored relative to baseline activity (prior to induction of hyperthermia).

For analysis of calcium transients, pre-processed traces were additionally high-pass filtered (0.01 Hz cutoff, 2^nd^ order Butterworth) to attenuate low-frequency drifts. Transients were programmatically determined using “findpeaks” in MATLAB 2023b using a threshold of a minimum of 1 peak prominence and a maximum width of 8 seconds. Transient width was defined at 25% of full peak prominence. Transient amplitudes were calculated from peak-to-trough by quantifying the local minimum within a 1-second window and subtracting it from the peak value. Inter-transient interval was calculated as the duration of time between one peak and the following peak. Transient frequency was calculated as the total number of peaks detected within each period (i.e., baseline, early heating, late heating, seizure) divided by the total duration of that period.

MATLAB 2023b was used to generate all figures that included representation of EEG and/or fiber photometry signals (GitHub repository: https://github.com/mattis-laboratory/ppn-chat-hyperthermia). GraphPad Prism was used to perform statistical analyses of the photometry signal and to generate summary plots.

### Immunohistochemistry

Animals were transcardially perfused with 4% paraformaldehyde (PFA) in 1x phosphate-buffered saline (PBS) for tissue analysis. Brains were post-fixed in 4% PFA for 24 hours and then transferred to 30% sucrose in PBS to equilibrate for at least 48 hours before slicing. 40 µm sections were collected using a frozen microtome (Leica Biosystems) and stored in cryoprotectant (25% glycerol, 30% ethylene glycol in PBS, adjusted to pH 6.7) at −20°C.

For immunostaining, brain slices were washed in PBS (three 10-minute washes) and then blocked and permeabilized with 3% normal donkey serum (NDS) in 0.3% Triton-X in PBS for 30 minutes. Slices were incubated in fresh blocking solution (3% NDS) in 0.3% Triton-X-100 in PBS) containing the primary antibody, rabbit polyclonal ChAT (1:1000; Thermo Fisher Scientific, Cat# PA5-29653, RRID:AB_2547128) for 48 hours at room temperature. The following primary antibodies were incubated for 24 hours at 4°C: mouse anti-GFAP (1:500; Sigma-Aldrich, Cat#G3893, RRID: AB_477010), and guinea pig anti-Iba1 (1:500; Synaptic Systems, Cat#HS-234 308, RRID:AB_2924932). Following incubation, sections were washed three times in PBS for 10 minutes each. The sections were then incubated for 3 hours at room temperature in blocking solution containing secondary antibody, diluted 1:500 (either Alexa Fluor 594 AffiniPure Donkey Anti-Rabbit, Jackson ImmunoResearch Labs, Cat# 711-585-152, RRID:AB_2340621, or goat anti-guinea pig Alexa Fluor 488 (Thermo Fisher Scientific, Cat#A-11073, RRID:AB_2534117). The tissue was washed in PBS three times for 10 minutes each, then stained with DAPI (1:50,000; Thermo Fisher Scientific, Cat# D1306) for 15 minutes. Sections were subsequently mounted and cover-slipped using PVA-DABCO mounting medium (MilliporeSigma, Cat# 10981).

For FluoroJade C staining, sections were mounted on slides and dried overnight. Then, FluoroJade C staining was carried out following the manufacturer’s instructions (Biosensis, TR-100-FJ), and slides were cover-slipped with DPX mounting media (Sigma-Aldrich, Cat#06522).

### Stereology and colocalization quantification

Cholinergic neuron number was quantified via the optical fractionator stereological technique (West et al., 1991). First, using a Nikon Fluorescent microscope (Nikon Eclipse Ni-E), a blinded investigator collected z-stacks with a grid step of 207 µm:203 µm (x:y) with a ×100 oil objective (Nikon) using systematic random sampling. Each z-stack spanned the entire thickness of the tissue in 1 µm steps. Next, cell counting was performed with the StereoInvestigator software (MBF Biosciences) using the Optical Fractionator probe and the nucleus as the counted object. Cell volume was measured using the Nucleator probe concurrently with the Optical Fractionator probe to ensure that only sampled cells were measured. 4 rays were randomly oriented on the cell and outlined the outside border of the cell and the outside border of the nucleus to obtain the whole cell volume, cytoplasm volume, and nucleus volume. Cell diameter was manually calculated from the volume estimates, assuming a spherical cell. All cell counts were performed by two independent researchers, both of whom were blinded to the condition. For cases in which the initial estimates differed by more than 15%, both researchers re-counted, again blinded to the condition and the direction of the initial count disparity.

For quantification of colocalization between ChAT and tdTomato expression, 2-3 sections were imaged per brain, capturing anatomical subregions across the cPPN. The region was first identified at low magnification, and tiled imaging was performed to ensure complete coverage of the area. Cell counting was performed live using a 60x oil-immersion objective (field of view: 222 µm x 221 µm), scanning through the Z-plane. Cells touching the field-of-view border were excluded from the count. Statistical analyses were performed, and graphs were generated using GraphPad Prism, as detailed above.

## Results

### REM sleep entry is impaired in DS mice following hyperthermic seizure induction

We began by performing polysomnography recordings of DS mice versus WT age and sex-matched littermate controls to characterize the combined impacts of genotype and recent seizure history on REM sleep. During a baseline (pre-seizure induction) 6-hour polysomnography recording, we detected no overall significant genotypic differences in REM sleep (**Figure 1; Figure S1**). However, we anecdotally noted that one DS mouse exhibited two spontaneous seizures during habituation (i.e., the day prior to baseline), and that this mouse subsequently had less REM sleep on the baseline day compared with the other mice in the cohort.

**Figure 1.**
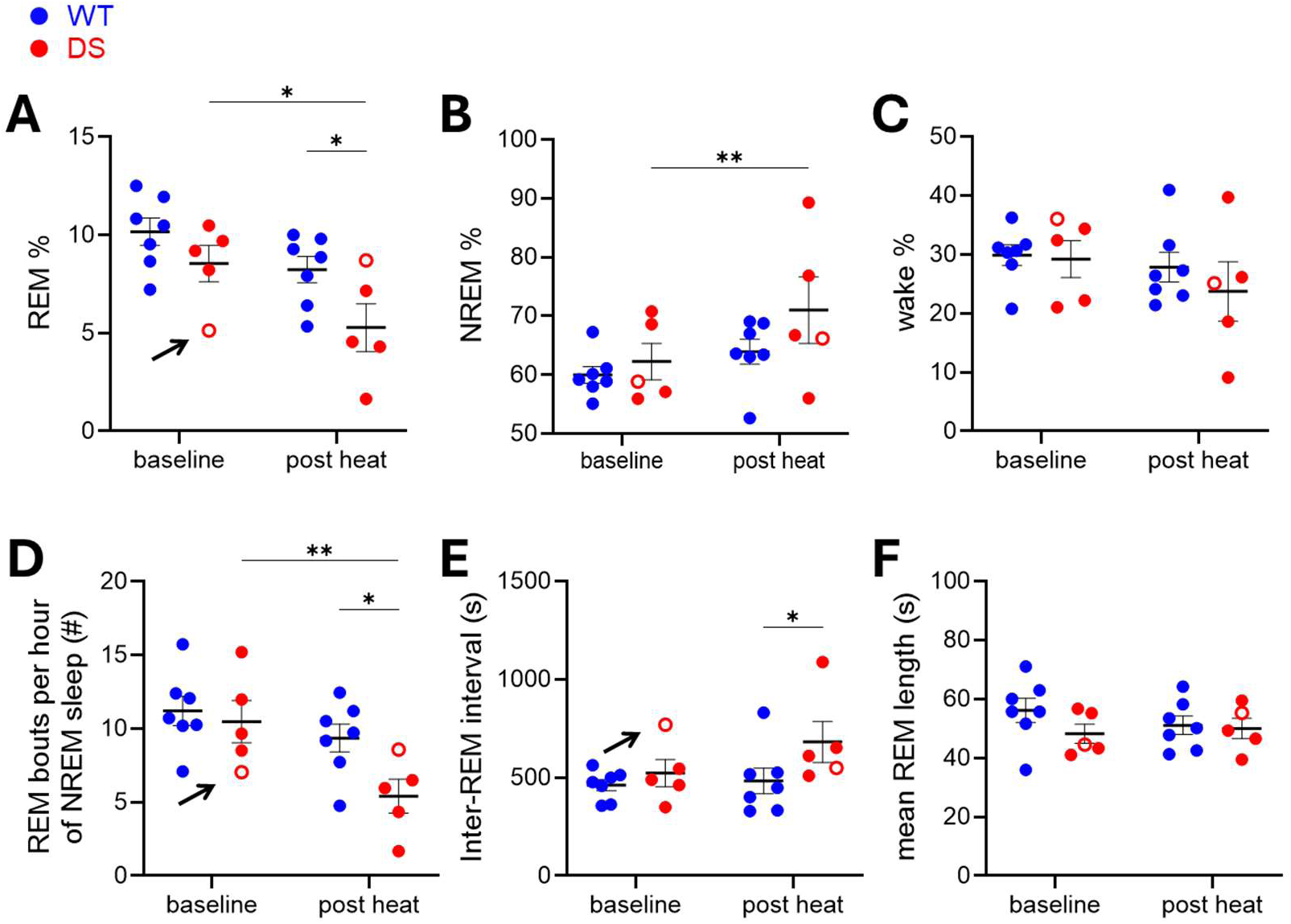
REM sleep entry is impaired in DS mice following hyperthermic seizure induction. **(A)** REM % was lower in post seizure DS mice (5.3 ± 1.2 %) compared to the same mice at baseline (8.5 ± 0.9 %; p = 0.04) and to WT mice after hyperthermia (8.2 ± 0.7 %; p = 0.02). **(B)** NREM % was higher in post seizure DS mice (71.0 ± 5.6 %) compared to the same mice at baseline (62.3 ± 3.1 %; p = 0.0099). WT mice after hyperthermia trended lower (64.0 ± 2.0 %; p = 0.12) **(C)** Wake % at baseline and post hyperthermia was similar across groups. **(D)** There were fewer REM bouts per hour of NREM sleep in post seizure DS mice (5.4 ± 1.2 bouts) compared to the same mice at baseline (10.5 ± 1.4 bouts; p = 0.008) and to WT mice after hyperthermia (9.4 ± 1.0 bouts; p = 0.02). **(E)** Inter-REM interval was higher in post seizure DS mice (682 ± 104 seconds) compared to WT mice after hyperthermia (483 ± 65 seconds; p = 0.05). DS mice at baseline trended lower (523 ± 69 seconds; p = 0.15). **(F)** Mean REM length was similar across groups. n = 7 mice for WT and n = 5 mice for DS. Open circles indicate data from the DS mouse that exhibited spontaneous seizures prior to the baseline recording; arrows indicate outlier values for that mouse. Pair-wise comparisons across genotypes and conditions were performed using Fisher’s Least Significant Difference (LSD) test. * p < 0.05, ** p < 0.01.

The following day, we exposed all mice to hyperthermia (to induce seizures in the DS mice) and then recorded a second 6-hour polysomnography recording (“post heat”), time-matched to the baseline recording to avoid circadian effects. During this second recording, DS mice exhibited a significant decrease in REM % relative to WT controls (**Figure 1A**), corresponding with an increase in non-REM (NREM) sleep but not wake (**Figure 1B-C; Table 1**). DS mice also exhibited a reduction in REM power spectral density (PSD), most pronounced within the theta range (**Figure S1**). We further found a reduced number of REM bouts per hour of NREM sleep (**Figure 1D**) and an increase in the inter-REM interval (**Figure 1E**), but no change in the mean REM bout duration (**Figure 1F**). Thus, the reduction in REM % following evoked seizures was specifically attributable to impaired REM entry.

### Hyperthermia-evoked seizures result in a significant increase in calcium signaling within DS PPN cholinergic neurons

The brainstem pedunculopontine nucleus (PPN) is a primary source of cholinergic inputs to the thalamus and brainstem that exerts regulatory control over REM sleep (Luo et al., 2023). Specifically, the PPN regulates REM entry, as opposed to REM sleep duration (Van Dort et al., 2015). Given our finding that recent hyperthermic seizure induction impaired REM entry, we hypothesized that seizures acutely impact the PPN in the DS model. We tested this hypothesis using photometry to record from PPN cholinergic neurons *in vivo* during seizures.

We crossed *Scn1a^+/-^*and *Chat^tm2(cre)Lowl/^J* (ChAT-Cre) mice (Rossi et al., 2011), stereotaxically introduced a Cre-dependent calcium indicator (GCaMP8s) into the PPN, and implanted an optical fiber over the PPN in addition to cortical EEG screws with a reference over the cerebellum (**Figure 2A-D**). Having confirmed that our recorded GCaMP signal corresponded with transitions into and out of REM sleep (**Figure S2**), as expected for PPN cholinergic neurons (Rye, 1997) we used a custom experimental rig to simultaneously acquire fiber photometry, EEG, and body temperature data during acute hyperthermic seizure induction.

**Figure 2.**
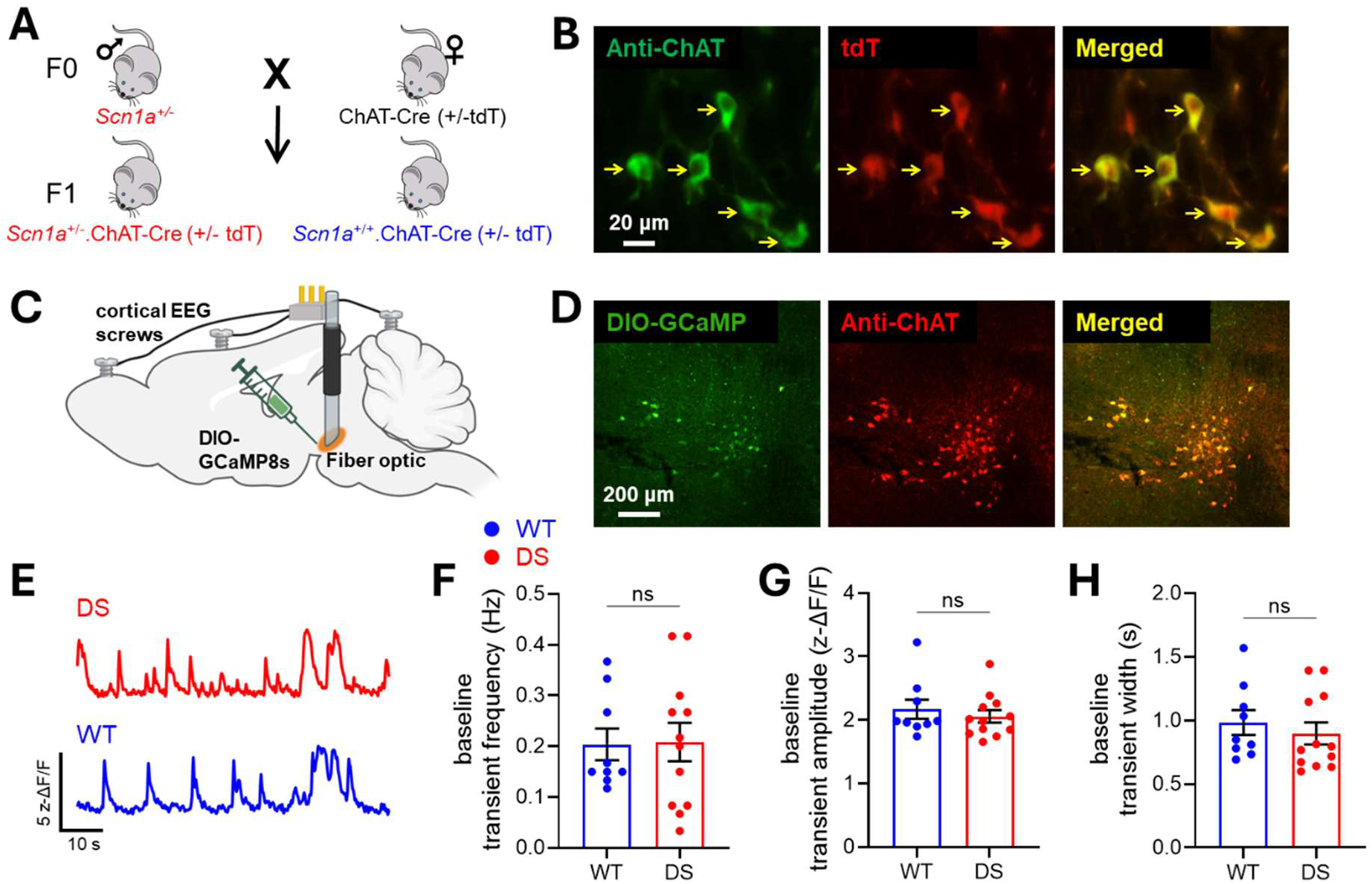
DS and WT PPN cholinergic neurons exhibit similar interictal population-level activity *in vivo*. **(A)** Breeding scheme to generate DS-ChAT mice and controls. **(B)** Representative fluorescent image of PPN neurons, indicating high overlap between ChAT immunostaining (green) and tdT expression (red). Yellow arrows indicate colocalized cells. **(C)** Schematic illustrating GCaMP injection and fiber optic placement in PPN, in combination with cortical skull screws for EEG recording. **(D)** Representative histology demonstrating robust GCaMP expression (green) within PPN neurons that immunostain for ChAT (red). **(E)** Representative spontaneous interictal calcium transients recorded from DS (red) and WT (blue) PPN cholinergic cells. **(F)** Baseline transient frequency was similar across genotypes (DS 0.21 ± 0.05 Hz; WT 0.20 ± 0.04 Hz; p = 0.83). **(G)** Baseline transient amplitude was similar across genotypes (DS 2.06 ± 0.14 z- ΔF/F; WT 2.17 ± 0.18 z-ΔF/F; p = 0.52). **(H)** Baseline transient width was similar across genotypes (DS 0.90 ± 0.12 s; WT 0.98 ± 0.12 s; p = 0.41). For panels F-H, statistical comparisons between genotypes were performed using a nested t-test on baseline data from n = 12 recordings (6 mice) for DS and n = 9 recordings (6 mice) for WT.

We recorded a total of 13 hyperthermia sessions (12 of which elicited seizures) across 6 DS-ChAT mice. As a control for the non-seizure-related effects of hyperthermia, we recorded 11 sessions from 7 WT littermate controls; notably, 2 sessions (from the same mouse) elicited unanticipated seizures / epileptiform activity and were therefore excluded from population analyses (**Figures S3-4**). To characterize the peri-ictal dynamics of the fluorescent response in PPN, we divided the sessions that contained seizures into four time periods of interest: baseline (pre-heating), early heating, late heating, and seizure; equivalent time periods were defined for WT sessions (without seizures) matched by heating durations.

We first analyzed the baseline period to detect any genotype differences in spontaneous activity *in vivo*. We observed no significant genotypic difference in the frequency, amplitude, or width of spontaneous calcium transients recorded (**Figure 2E-H**). To ensure that z-scoring was not obfuscating genotypic differences in signal amplitudes, we additionally compared amplitudes from non-z-scored ΔF/F traces and again found no significant differences between genotypes.

We next analyzed the GCaMP signal obtained during seizures (or equivalent heating period), z-scored relative to baseline values. We observed a significant increase in the signal from PPN cholinergic cells during seizures in DS-ChAT mice, but not in the equivalent WT traces (**Figure 3A-C**). This sharp rise in fluorescence coincided most closely with the earliest electrographic change around the onset of the seizure (EEC; **Figure 3D**) that continued into the unequivocal electrographic onset of the seizure (UEO; **Figure 3E**) and declined abruptly at seizure offset (OFF; **Figure 3H**). We quantified the area under the curve (AUC) of each triggered average (**Figure 3G-I**), re-demonstrating the significant seizure-associated increase in fluorescent signal.

**Figure 3.**
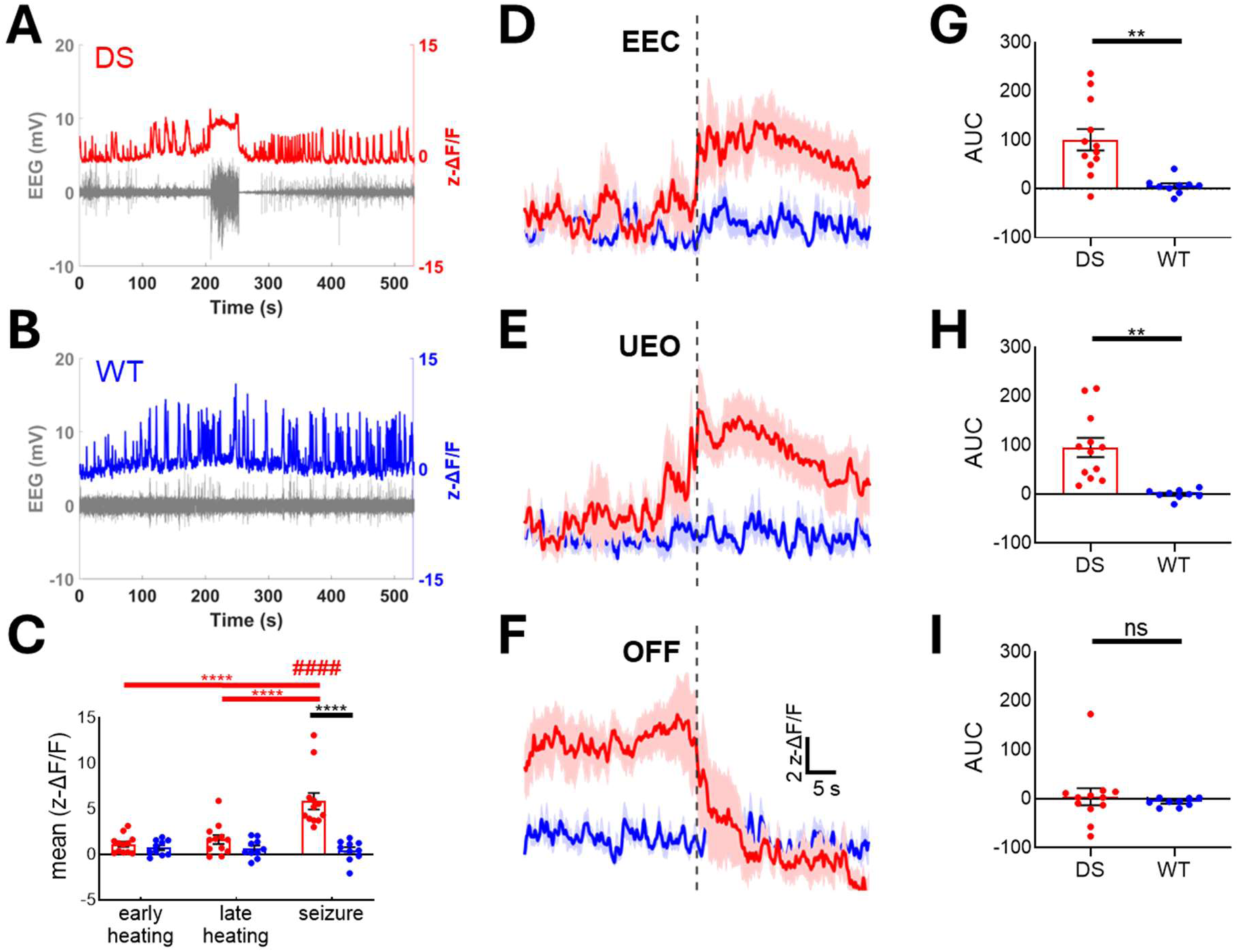
Hyperthermia-evoked seizures trigger an acute increase in GCaMP signal in DS cholinergic neurons. **(A)** Example EEG trace (mV; gray) overlaid with fiber photometry signal (z-ΔF/F; red) recorded from a DS-ChAT mouse before, during, and after a hyperthermia-evoked seizure. **(B)** Example EEG trace (mV; gray) overlaid with synchronized fiber photometry signal (z-ΔF/F; blue) recorded from a WT control mouse that was subjected to hyperthermia but did not exhibit a seizure. **(C)** Mean z-ΔF/F values recorded during the early heating, late heating, and seizure (or seizure-equivalent) portions of the recording. Photometry signals were z-scored relative to the baseline (pre-heating) recording period. 2-way ANOVA demonstrated a significant effect of time-period (p = 0.0003), genotype (p = 0.03), and an interaction between time-period and genotype (p < 0.0001). Mean DS fluorescence during the seizure was significantly greater versus all other time periods within those mice (****; p < 0.0001), and versus the equivalent time-period in WT mice (####; p < 0.0001). **(D)** Triggered average of fluorescence signal aligned to the EEC for DS-ChAT mice (red), and to the equivalent timepoint in WT mice (blue). Dashed line denotes time of EEC or equivalent timepoint. Solid lines and shading represent mean and SEM, respectively. **(E)** Triggered average aligned to the UEO / equivalent time point. **(F)** Triggered average aligned to OFF / equivalent time point. **(G)** Area under the curve (AUC) of EEC-triggered averages (DS 101 ± 22; WT 6.13 ± 5.50; p = 0.005. **(H)** AUC for UEO-triggered averages (DS 95.42 ± 19.65; WT 0.025 ± 3.27; p = 0.0021). **(I)** AUC for OFF-triggered averages (DS 4.36 ± 17.46; WT −6.89 ± 2.83; p = 0.50). n = 9 recordings (6 mice) for WT and n = 12 recordings (6 mice) for DS. *p < 0.05, **p < 0.01, ***p < 0.001, ****p < 0.0001. Triggered averages were calculated across the first 30 seconds after EEC/UEO/OFF for DS or after equivalent timepoints for WT. For panel C, pair-wise statistical comparisons were performed using Šídák’s multiple comparisons test. For panels G-I, significance was determined using nested t-tests.

### DS PPN cholinergic neurons are reduced in number and exhibit morphologic changes following repeated hyperthermia-evoked seizures

Previous work has identified reduced PPN neuron numbers (Hayashi et al., 2012), PPN atrophy (Cho et al., 2016), and reduced free cerebrospinal fluid free choline levels (Pranzatelli et al., 1998) in patients with epilepsy. We therefore hypothesized that seizures in DS mice would have lasting effects in the PPN beyond the acute ictal period. As induction of seizures within an early developmental time window is known to exacerbate the phenotype of DS model mice (Hawkins et al., 2017, 2019; Guo et al., 2024; Zhou et al., 2024; Yip et al., 2025), we subjected mice to repeated hyperthermia-evoked seizures at P19, P21, and P23 (“3x heat” group), harvesting their brains 2-3 hours after the third heating session. To disambiguate the effects of genotype (in the absence of induced seizures), we ran a second group in parallel (“no heat”) in which no hyperthermia was administered. Both groups included WT littermate control mice that underwent the same procedures but did not exhibit behavioral seizures.

We observed overall reduced ChAT immunostaining in sections from DS mice that had been subjected to repeated hyperthermia (**Figure 4A**). To precisely quantify PPN cholinergic neuron number across experimental groups, we employed the optical fractionator stereology approach (**Figure 4B**). We focused on the caudal PPN (cPPN) as it is more highly enriched for cholinergic neurons (Mena-Segovia et al., 2009), and tiled the region with a series of high magnification z-stacks. Using StereoInvestigator, we counted the number of cholinergic neurons within the dissector and extrapolated an estimate of the total cPPN cholinergic neuron number for each animal. The dissector was set to 75 x 75 µm with a thickness of 14.5-20 µm without guard zones (**Table 1**). To ensure that the estimate of cell number was accurate, we established sampling parameters that achieved a coefficient of error (Gundersen m = 1) that was consistently less than 0.1, as is standard in the field (Gundersen et al., 1999) (**Table 1; Figure S5A-B**). We confirmed that the cell number count was not sensitive to sections counted – i.e., that we had avoided under-sampling (**Figure S5C**).

**Figure 4.**
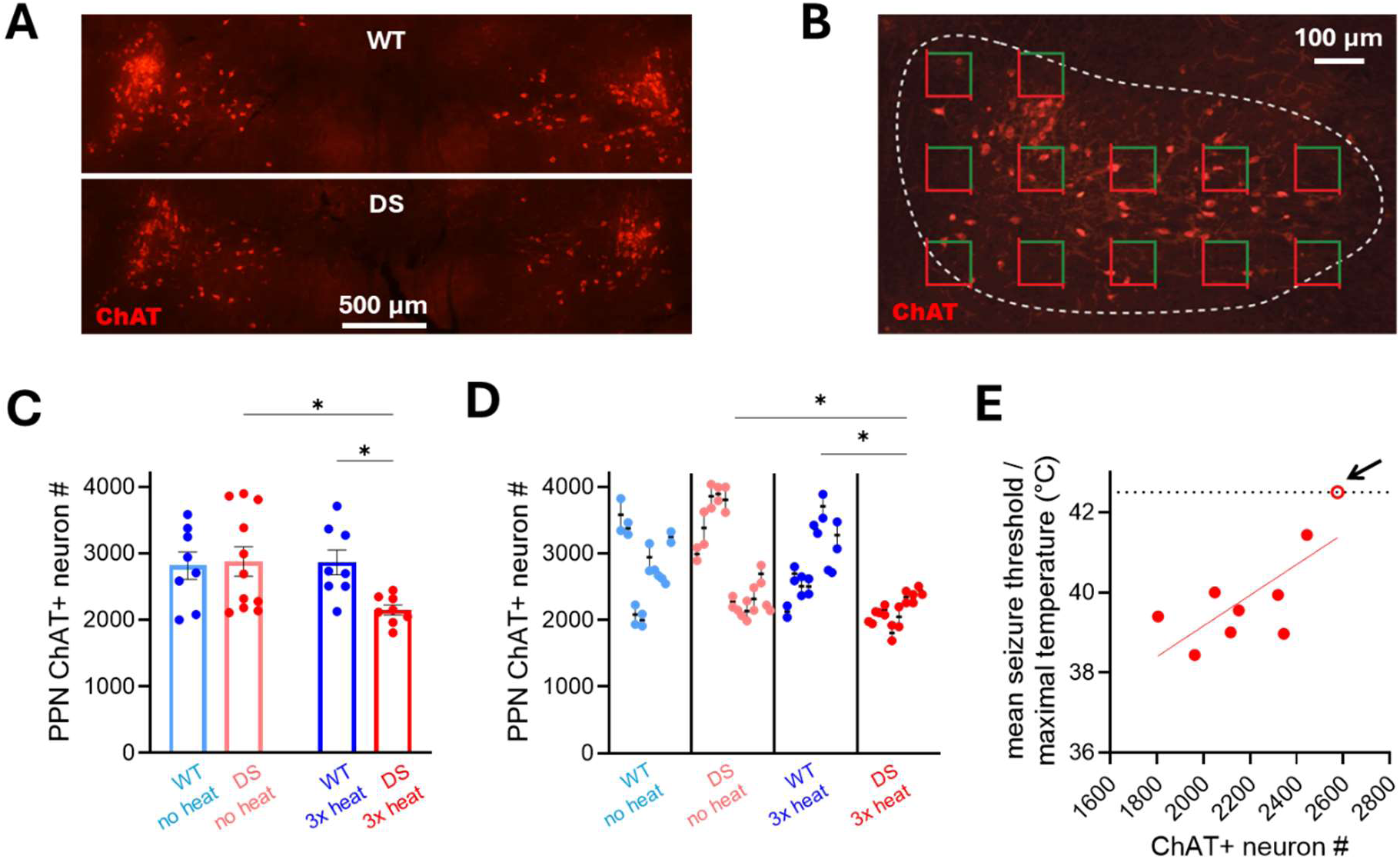
DS mice subjected to repeated hyperthermia-evoked seizures have fewer caudal PPN cholinergic neurons. **(A)** Representative images of the cPPN from a 3x heat-treated WT mouse (above) and DS mouse (below), showing reduced ChAT immunoreactivity in the DS mouse. **(B)** Illustration of the Optical Fractionator approach using systematic random sampling for unbiased quantification of cholinergic neurons (red) within the cPPN (dashed outline). The dissector is shown as squares with red and green sides. **(C)** Estimates of cPPN cholinergic neuron numbers for WT and DS mice across no heat and 3x heat groups. Counts (mean ± SEM) were similar across WT no heat (baby blue, 2815 ± 206), DS no heat (pink, 2876 ± 222), and WT 3x heat (2864 ± 187) groups but the DS 3x heat group had a neuron count (2150 ± 75) that was significantly lower than both the DS no heat (p = 0.018) and the WT 3x heat (p = 0.010) groups. Pair-wise comparisons were performed using Fisher’s Least Significant Difference (LSD) test. **(D)** Stereological counts from each of two blinded experimenters. Nested multiple comparisons revealed significant differences between DS no heat and 3x heat mice (p = 0.019) and between WT and DS 3x heat mice (p = 0.019). **(E)** Seizure threshold (averaged across all heating sessions for each mouse) was positively correlated with ChAT neuron estimate (R^2^ = 0.53, p = 0.026, linear regression). One DS mouse (open circle; arrow) did not exhibit a seizure in any heating session and had a ChAT neuron estimate (2575 cells) higher than any other DS mice in the 3x heat group. Note that for this mouse, the maximal temperature recorded (42.5 °C) was used for analysis. For the no heat group, n = 8 mice (WT) and 11 mice (DS); for the 3x heat group, n = 8 mice (WT) and 8 mice (DS).

In the no heat group, our estimates were in line with previous literature (Mena-Segovia et al., 2009; Luquin et al., 2018; Soares et al., 2018), and we observed no genotypic difference in cPPN cholinergic neuron number (**Figure 4C**). However, in the 3x heat group, DS mice had ∼25% fewer cPPN cholinergic neurons compared to both 3x heat WT controls and no heat DS mice. This finding was confirmed by two independent investigators, both of whom were blinded to condition (**Figure 4D; Table 1**). Within the 3x heat DS group, we found that ChAT+ neuron number had a significant positive correlation with mean seizure onset threshold (**Figure 2E**). Interestingly, a single DS mouse in the 3x heat group – whose genotype was confirmed by PCR x3, from both tail and brain tissue – failed to exhibit a behavioral seizure during any of the heating sessions; the number of cholinergic cells quantified from this mouse was higher than any of the other 3x heat DS mice (that did exhibit behavioral seizures). Overall, these data identify an association between hyperthermic seizure burden – distinct from the DS genotype – and cPPN cholinergic neuron number.

We considered that the decrease in cPPN cholinergic neurons might be due to seizure-induced cell death. To test this hypothesis, we used FluoroJade C (FJC), a marker that labels degenerating neurons independent of the mechanism of cell death (Gu et al., 2012). We also tested for inflammation within the PPN (Elson et al., 2016), staining for ionized calcium-binding adapter molecule 1 (Iba1) and glial fibrillary acidic protein (GFAP) to detect microglial and astrocytic activation, respectively (Ito et al., 2001; Brahmachari et al., 2006). Hippocampal slices from kainic acid-treated mice were stained concurrently as a positive control (Bendotti et al., 2000; Ding et al., 2000; Zhang and Zhu, 2011). No notable differences were detected in the PPN following hyperthermia-evoked seizures in DS mice relative to WT controls exposed to hyperthermia (**Figure 5A**).

**Figure 5.**
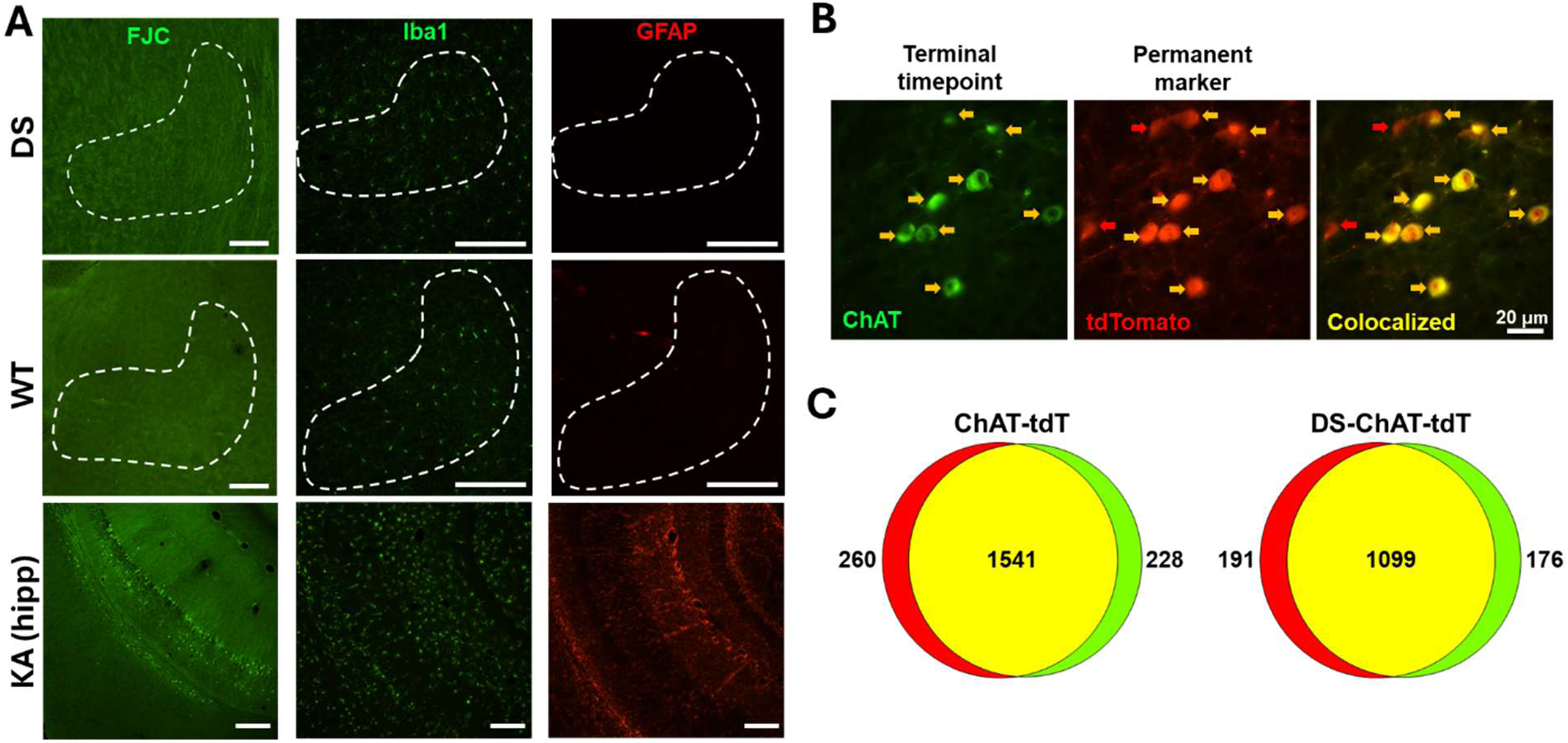
Repeated hyperthermia-evoked seizures do not induce cell death or a loss of ChAT expression within PPN cholinergic neurons. **(A)** FJC staining (left column; green) was not observed in the PPN of heated DS (top row) or WT (middle row) mice but was seen in the ventral hippocampus of KA-treated mice (bottom row). Images are representative of results obtained in n = 5 DS mice, n = 6 WT mice, and n = 2 KA mice. No differences in PPN expression of the microglial marker Iba1 (center column; green) or astrocyte marker GFAP (right column; red) were observed between heated DS and WT mice, whereas strong staining was seen in the KA positive control. Similar results were obtained in n = 3 DS mice and n = 3 WT mice, n = 2 KA mice. All scale bars = 200 µm. **(B)** Representative images of PPN neurons that immunostain for ChAT (green) and/or express tdT (red). Yellow arrows indicate colocalized cells; red arrows indicate cells positive for tdT but not ChAT. **(C)** Quantification of tdT and ChAT co-expression in ChAT-tdT and DS-ChAT-tdT mice. Of tdT-expressing neurons, the percentage that immunostained for ChAT at the experimental endpoint was 85.6% for ChAT-tdT and 85.2% DS-ChAT-tdT mice. n = 2029 cells / 7 mice (ChAT-tdT) and 1466 cells / 5 mice (DS-ChAT-tdT); 2-3 sections per mouse.

Previous work has described an activity-dependent phenotypic shift of cholinergic cPPN neurons – in which they reversibly lose ChAT expression and gain GABAergic markers (Li et al., 2020) – in the context of exercise. To test whether seizures similarly induce a loss of ChAT expression, we crossed *Scn1a^+/-^* breeders with ChAT-Cre;tdT mice, in which tdT expression serves as a stable marker of a cell’s prior – not only current – cholinergic identity (given that Cre-mediated deletion of the STOP cassette results in irreversible tdT expression). We quantified co-localization of tdT expression (i.e., cells that were cholinergic at any timepoint) and ChAT immunostaining (i.e., cells that were cholinergic at the time of perfusion) (**Figure 5C**). We found no genotypic differences in the percentage of tdT+ cells that colocalized with ChAT immunostaining (1541/1801 tdT+ cells = 85.6% of tdT+ cells in WT (ChAT-tdT) and 1099/1290 tdT+ cells = 85.2% of tdT+ cells in DS-ChAT-tdT mice), indicating that repeated seizures do not shift ChAT expression in the cPPN (**Figure 5C**). These results were verified by a second independent quantification (**Figure S6**).

Finally, we observed that ChAT neurons appeared to be larger in the heated groups (**Figure 6A**). We therefore expanded our anatomic characterization within the PPN by measuring the volume of cPPN cholinergic neurons, using those cells that were sampled for stereology estimation, and found a significant effect of heating on ChAT neuron volume (**Figure 6B-C**) due to increases in both nucleus volume (**Figure 6D**) and cytoplasm volume (**Figure 6E**). Although hyperthermia impacted both genotypes, the relative effect was more pronounced for DS mice.

**Figure 6.**
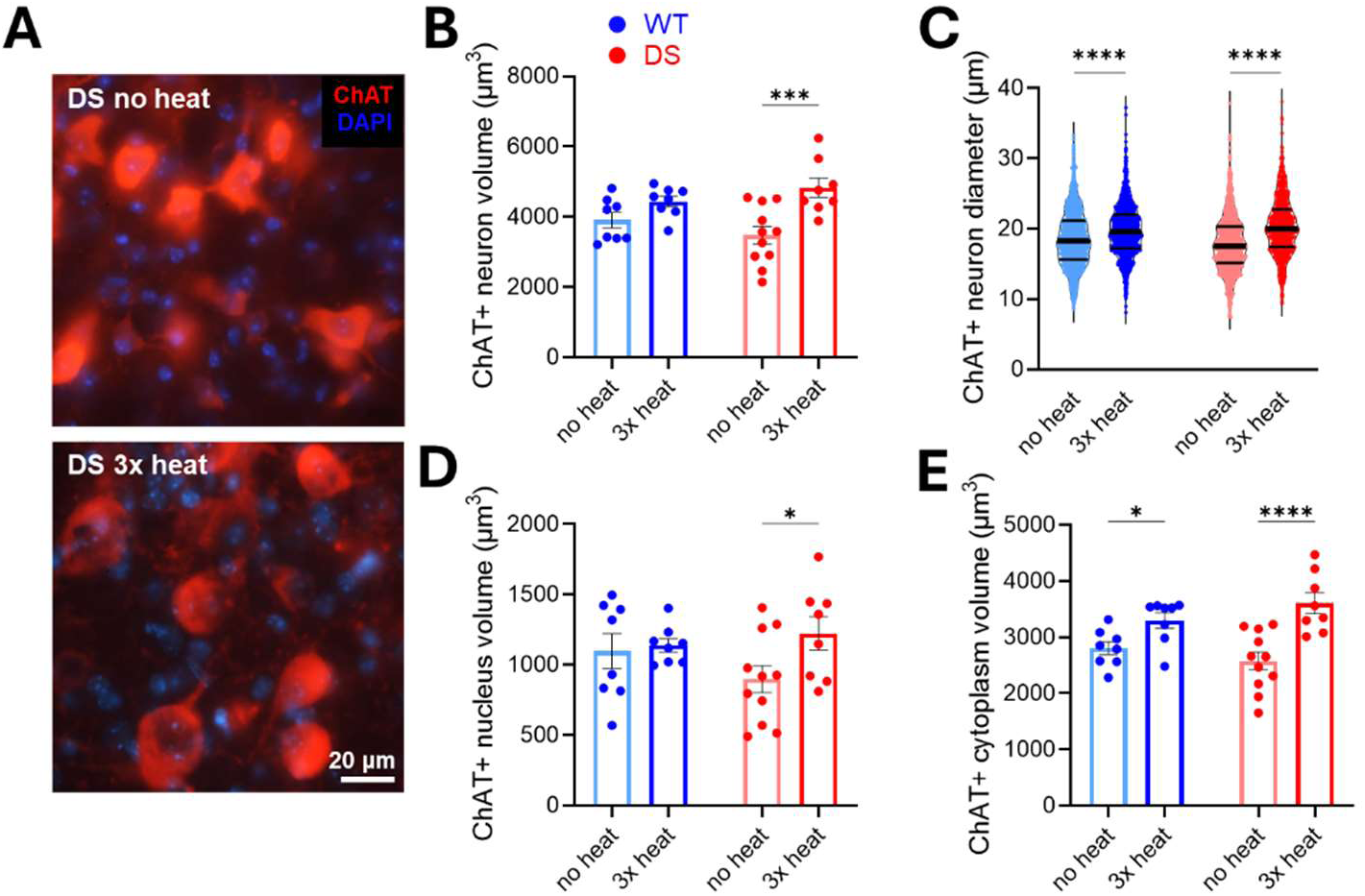
Mice subjected to repeated hyperthermia exhibit an increase in the PPN cholinergic neuronal size. **(A)** Representative images of cPPN cholinergic immunostaining (red) and nuclear DAPI expression (blue), illustrating larger cholinergic neuron volume in the 3x heat versus no heat DS groups. **(B)** ChAT total neuron volumes were averaged within each animal and compared across conditions. There was a significant overall effect of heating (p = 0.0004) and an increase between no heat DS (3475 ± 251 µm^3^) and 3x heat (4828 ± 273 µm^3^) DS neuron size (p = 0.0002). **(C)** Distribution of ChAT neuron diameters revealed a significant overall effect of heating as well as an interaction effect (p < 0.0001 for both). WT neurons increased in diameter by ∼6% (18.6 ± 0.1 µm no heat versus 19.8 ± 0.1 µm 3x heat; p < 0.0001), whereas the DS neurons increased diameter by ∼14% (17.8 ± 0.1 µm no heat versus 20.3 ± 0.1 µm 3x heat; p < 0.0001). **(D)** ChAT+ neuron nuclear volumes were averaged within each animal and compared across conditions. 3x heat DS volumes were increased relative to no heat DS (899 ± 96 µm no heat versus 1221 ± 119 µm 3x heat; p = 0.03). **(E)** ChAT neuron cytoplasm volumes were averaged within each animal and compared across conditions. There was an overall effect of heating (p < 0.0001), with a significant increase between no heat DS (2576 ± 526 µm^3^) and 3x heat DS (3606 ± 533 µm^3^; p < 0.0001), as well as no heat WT (2804 ± 329 µm^3^) and 3x heat WT (3300 ± 390 µm^3^; p = 0.04). For the no heat group, n = 8 mice (WT) and 11 mice (DS); for the 3x heat group, n = 8 mice (WT) and 8 mice (DS). All overall effects were obtained by two-way ANOVA. Pair-wise comparisons were performed using Fisher’s Least Significant Difference (LSD) test. In panels B, D, and E, values were averaged across all cells per animal (n = mouse), and data are presented as mean ± SEM. In panel C, n = neuron (1059-1444 per group), and data are presented as a violin plot in which lines indicate median and quartiles. * p < 0.05, *** p < 0.001, **** p < 0.0001.

## Discussion

### Impaired REM entry following hyperthermic seizure induction in DS mice

REM sleep impairment occurs following spontaneous seizures in patients with a range of epilepsy types (Bazil et al., 2000; Kilgore-Gomez et al., 2024) and has been demonstrated in a preclinical model of temporal lobe epilepsy (Nair et al., 2022). However, prior characterizations of sleep architecture in the DS mouse found reduced theta-range power during REM but identified no differences in either REM % (Kalume et al., 2015) or REM bout duration (Sanchez et al., 2019). Notably, however, these studies were performed without regard to seizure recency. Although we observed no significant differences in REM sleep between DS and WT mice at baseline (prior to seizure induction), in the hours following hyperthermic seizure induction, we detected a significant reduction in REM %, attributable to impaired REM entry (**Figure 1**).

What is the potential clinical relevance of this finding? Patients with DS have significantly disrupted sleep (Licheni et al., 2018), with frequent reports of nighttime awakenings and daytime sleepiness (Schoonjans et al., 2019). Although causality is certain to be complex in this severe developmental and epileptic encephalopathy, survey data collected from hundreds of caregivers support a strong positive correlation between seizure burden and the number and severity of comorbid symptoms (Lagae et al., 2018; Makiello et al., 2023). Conversely, patients with DS exhibited improved executive functioning following seizure reduction (Bishop et al., 2021). If our pre-clinical observation holds in the clinical DS population, seizure-induced impairment of REM sleep may be one mechanism by which a high seizure burden can incur lasting cognitive effects.

### Interictal and ictal activity of PPN cholinergic neurons in DS mice, as measured by *in vivo* fiber photometry

As the PPN is known to regulate REM sleep entry (Shiromani and McGinty, 1986; Webster and Jones, 1988; Van Dort et al., 2015), we identified it as a candidate region to investigate post-seizure REM sleep disruption in DS mice.

During the interictal period, we found no evidence that *Scn1a* haploinsufficiency overtly alters the endogenous activity of PPN cholinergic neurons (**Figure 2**), although it has been reported that at least one third of PPN cholinergic neurons express Nav1.1 (Papale et al., 2013). Our recent work notably demonstrated preserved functional properties of cholinergic neurons within the medial septum of DS mice (Zhu et al., 2025), in which ∼90% of ChAT neurons express Nav1.1 (Bender et al., 2016). It is possible that cholinergic neurons do not rely strongly on Nav1.1 for action potential generation, or that they can compensate for reduced Nav1.1 expression via upregulation of other sodium channel subtypes, as reported for other neuronal populations (Yu et al., 2006; Catterall et al., 2010; Chopra and Isom, 2014). Prior work (in neocortical inhibitory neurons) demonstrated that Nav1.1 haploinsufficiency significantly impairs action potential propagation (Kaneko et al., 2022), thus it remains possible that cholinergic neurotransmission to downstream target regions is disrupted in DS.

During the ictal period, we identified a dramatic genotypic divergence in PPN cholinergic neuron activity, with a highly significant increase in GCaMP signal coinciding with early electrographic seizure onset in DS mice but not in WT controls exposed to equivalent hyperthermia (**Figure 3**). Previous studies have investigated PPN activity during seizures, but with varying results. During evoked limbic seizures – triggered via electrical hippocampal stimulation in anesthetized rats and awake behaving mice – juxtacellular in vivo recordings in the PPN showed an overall decrease in cholinergic neuronal firing (Motelow et al., 2015; Liu et al., 2025). In contrast, during flurothyl-evoked generalized seizures in WT rats, the PPN displayed increased activity, as determined by a 2-deoxyglucose autoradiography mapping technique (Velíšková et al., 2005). Our data align with the latter study, supporting a model in which the PPN is activated by generalized seizures but inhibited by focal onset seizures (at least those originating in the temporal lobe). Alternatively, the sustained fluorescence peak that we observed may represent depolarization block, in which case this increased signal would not reflect sustained action potential firing. Future studies that directly measure downstream acetylcholine release during seizures will be needed to clarify this point.

What might be the consequences of this acute ictal disruption of PPN cholinergic networks? Given the PPN’s role in arousal and brain state regulation (Mena-Segovia et al., 2008; Benarroch, 2013; Collazo et al., 2019), disruption of its physiologic activity may contribute to impaired consciousness during seizures. The PPN also innervates multiple brainstem nuclei involved in autonomic regulation (Mena-Segovia et al., 2008; Martinez-Gonzalez et al., 2014; Mena-Segovia and Bolam, 2017), so alteration of PPN efferent firing may contribute to seizure-associated autonomic dysfunction.

### Impact of repeated seizures on PPN cholinergic neurons in DS mice

To build upon our acute observations, we interrogated the impact of repeated seizures on the PPN on a longer timescale. We observed that DS mice subjected to repeated hyperthermia-induced seizures exhibited a ∼25% reduction in cPPN cholinergic neuron number (**Figure 4**). These results can again not be attributed to genotype alone, as these were no different between DS and WT mice that were not subjected to heating. Because spontaneous seizures in DS mice begin by ∼P14 (Han et al., 2020), the non-heated DS group may have exhibited seizures, which we did not measure or quantify, prior to the experimental endpoint. However, induced hyperthermia seizures have been well-established to exacerbate disease severity in DS mice (Hawkins et al., 2017, 2019; Guo et al., 2024; Zhou et al., 2024; Yip et al., 2025), including increased spontaneous seizure frequency. Given this knowledge, we interpret our data to suggest a dose-dependent response, such that only the heated group experienced a sufficient seizure burden to trigger the observed changes within the PPN. However, an alternative explanation – which we cannot exclude – is that hyperthermia itself triggered these changes within PPN, with a more profound impact on DS mice compared to WT counterparts. The higher neuron number in the n = 1 DS mouse that repeatedly failed to exhibit seizures in response to hyperthermia (**Figure 4E**) argues otherwise, but this outlier only provides anecdotal evidence. Future studies could be designed to dissociate the heat and seizures exposures, e.g., by treating DS mice with anti-seizure medications to prevent seizures despite hyperthermia exposure or triggering seizures via a chemoconvulsant instead of hyperthermia to clarify their relative contributions.

Our work fits within a larger literature from clinical and preclinical studies suggesting that cholinergic networks, including the PPN, are chronically disrupted in epilepsy. Postmortem analyses of patients with a variety of refractory epilepsy disorders (West Syndrome, dentatorubral-pallidoluysian atrophy, and Lennox-Gastaut Syndrome) revealed a decrease in the number and percentage of cholinergic PPN neurons (Hayashi et al., 2012). Patients with epilepsy have altered choline levels in cerebrospinal fluid choline levels (Pranzatelli et al., 1998; Oates et al., 2023) as well as altered expression of acetylcholine receptors (Sun et al., 2021). Patients with sleep-predominant seizures displayed a volume reduction of the PPN, as measured by magnetic resonance imaging (MRI) (Cho et al., 2016). Finally, functional MRI (fMRI) data from patients with temporal lobe epilepsy revealed decreased connectivity between the PPN and its cortical and subcortical targets (Englot et al., 2009, 2018), which recovered in patients who achieved postoperative seizure freedom (González et al., 2020).

By what mechanism might repeated hyperthermia-evoked seizures result in reduced cPPN cholinergic neuron numbers? While the most straightforward explanation is cell death, we surprisingly failed to detect evidence for this mechanism within the PPN (**Figure 5A**), although we cannot exclude the possibility that cell death occurred rapidly and concluded too early to be captured by our analysis. We tested the possibility of neurotransmitter switching, whereby individual neurons undergo a phenotypic shift and lose markers of their initial neurotransmitter expression, as previously reported in PPN (Li and Spitzer, 2020), but found no evidence to support this hypothesis in our experimental paradigm (**Figure 5B-C**). It is possible that higher cPPN cholinergic neuron numbers correlate with a lower probability of survival to the experimental endpoint – i.e., that our results are driven by survivorship bias, although the positive correlation between PPN cholinergic neuron number and seizure threshold (**Figure 4E**) suggests otherwise.

Finally, we observed that repeated hyperthermia-evoked seizures were associated with increased somatic volume of PPN cholinergic neurons (**Figure 6**). PPN cholinergic cell size is tightly regulated across development: a study in rats showed that PPN cholinergic neurons increase in size during early postnatal development, between P12 and P16, and subsequently start to decline at around P21 (Kobayashi et al., 2004). Thus, repeated seizures may disrupt the normal developmental trajectory of cholinergic neurons in the PPN, preventing the expected reduction in cell size. Alternatively, seizures (in combination with hyperthermia) may induce hypertrophy. Indeed, prior reports note cholinergic hypertrophy in response to a range of physiologic stimuli, including growth factor exposure, exercise, and aging (Higgins et al., 1989; Stroessner-Johnson et al., 1992; Hall et al., 2018; Kobayashi et al., 2004). Similarly, kainic acid-induced seizures in rats increased PPN (and neighboring lateral dorsal tegmental nucleus) ChAT neuron volume, accompanied by increased cholinergic varicosities in the parafascicular nucleus (Soares et al., 2018).

### Conclusions and future directions

This present work provides novel insights related to REM sleep disruption as well as PPN function in a DS mouse model. We demonstrate: (1) seizure-associated REM sleep disruption, (2) acute recruitment of PPN cholinergic neurons by seizures, and (3) subacute anatomic remodeling of PPN cholinergic neurons following repeated seizures. We propose a conceptual model whereby an acute ictal insult to the PPN triggers changes within PPN cholinergic neurons that cause impaired REM sleep entry. However, additional work remains to confirm the causality ascribed by this model. As a first step, it will be important to establish the generalizability of our findings across epilepsies and to establish whether only seizures that directly impact the PPN are associated with subsequent changes in PPN anatomy and sleep architecture.

The PPN has already been employed as a surgical target for deep-brain stimulation in patients with Parkinson’s disease (Garcia-Rill et al., 2019) and has been proposed as a therapeutic target in epilepsy (Jaseja, 2021). If seizure-associated REM sleep disruption is ultimately attributed to the PPN, future neuromodulatory strategies could be employed not only to mitigate seizures, but also to protect against seizure-associated sleep impairment that significantly worsens quality of life in epilepsy.

## Author contributions

Krystal Santiago-Colon, Chandni Rana, Julia A. Kravchenko, and Joanna Mattis contributed to project conceptualization. Krystal Santiago-Colon, Chandni Rana, and Joanna Mattis contributed to data curation. Krystal Santiago-Colon, Chandni Rana, Brandon Toth, Julia Kravchenko, Joseph Barden, and Joanna Mattis contributed to formal data analysis. Joanna Mattis contributed to funding acquisition. Krystal Santiago-Colon, Chandni Rana, Brandon Toth, Julia A. Kravchenko, Joseph Barden, Isabella VanHorn, Alexandra Jackson, Alexander Hollier, Rida Qureshi, Hudson Haberland, Juan Disla, Nagham Eldroubi, and Sophie Erb-Watson contributed to the research investigation process, that is, performing experiments and collecting data. Krystal Santiago-Colon, Chandni Rana, Brandon Toth, Julia A. Kravchenko, and Joanna Mattis contributed to the development and design of the methodology. Joanna Mattis was responsible for project administration and management. Chandni Rana, Brandon Toth, and Saif Siddiqui contributed to programming, software development, and implementation of analysis code. Joseph Barden and Joanna Mattis contributed to the provision of resources, including study materials, reagents, and animals. Joanna Mattis was responsible for oversight and leadership supervision. Lori Isom provided mentorship to Krystal Santiago-Colon. Krystal Santiago-Colon, Chandni Rana, Brandon Toth, Joseph Barden, and Joanna Mattis contributed to data visualization. Krystal Santiago-Colon, Chandni Rana, and Joanna Mattis wrote the original draft of the manuscript. All authors contributed to reviewing and editing the manuscript.

