## Supplementary figures and table for "The impact of seizures on REM sleep and the cholinergic pedunculopontine nucleus in a mouse model of Dravet Syndrome"

### Supplemental Figures

#### Figure S1. Power spectral densities at baseline and post heat.

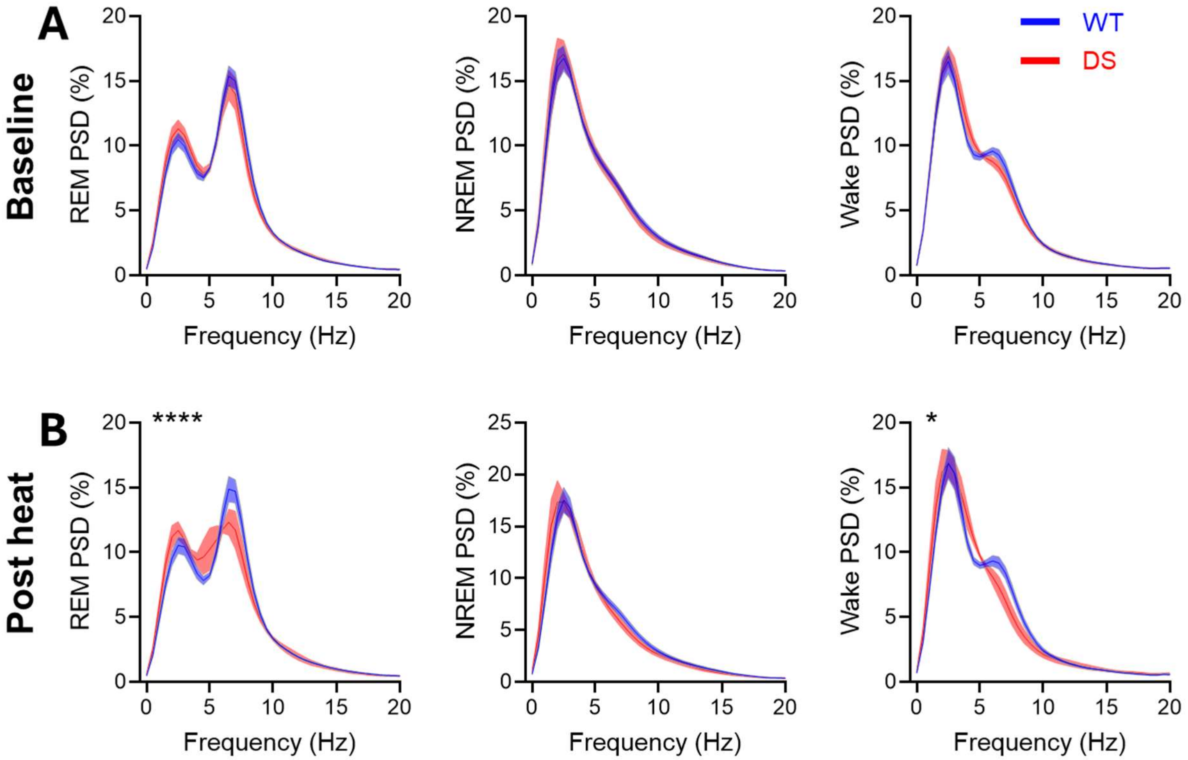


**(A)** Baseline REM (left), NREM (center), and wake (right) PSD plots, none of which showed significant genotype differences.

**(B)** Post heat PSD plots. REM PSD (left) displayed a highly significant interaction between genotype and frequency ($p < 0.0001$); power was reduced in DS mice at 6.5 Hz ($p = 0.009$), 7 Hz ($p = 0.0007$) and 7.5 Hz ($p = 0.016$) (Šidák multiple comparison test). NREM PSD (center) did not display genotype differences. Wake PSD (right) displayed a significant interaction between genotype and frequency ($p = 0.016$).

Stars indicate genotype/frequency interaction calculated from two-way repeated measures ANOVA. Note that all ANOVAs were highly significant for frequency ($p < 0.0001$).

#### Figure S2. Confirmation of *in vivo* targeting of PPN cholinergic cells based on response to sleep state transitions.

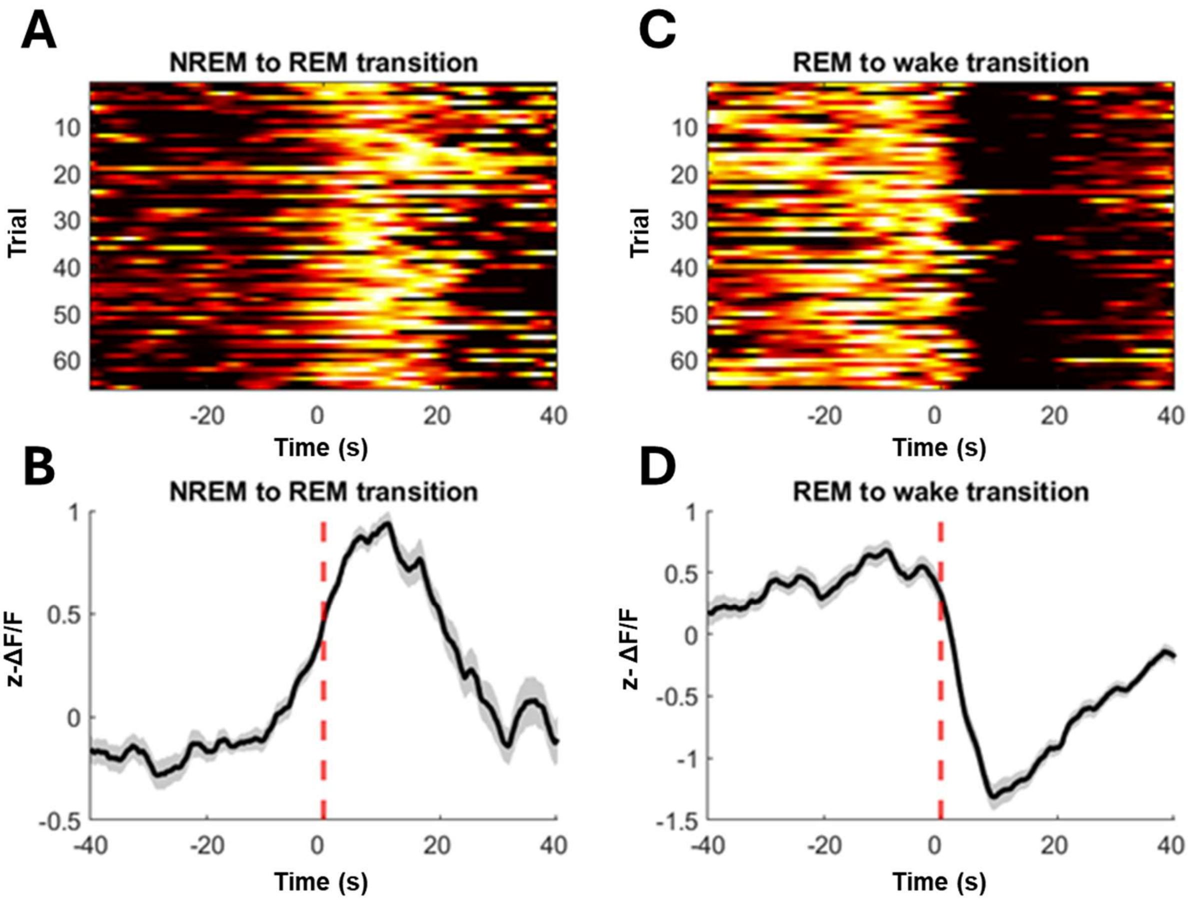


**(A)** Single trial transition-onset-triggered time courses of GCaMP fluorescence at NREM to REM transitions.

**(B)** Average GCaMP fluorescence aligned to NREM to REM transitions (dashed red line). Solid lines and shading indicate mean and SEM, respectively.

**(C)** Same as (A) for REM to wake transitions.

**(D)** Same as (B) for REM to wake transitions.

#### Figure S3. Summary of mice recorded with fiber photometry.

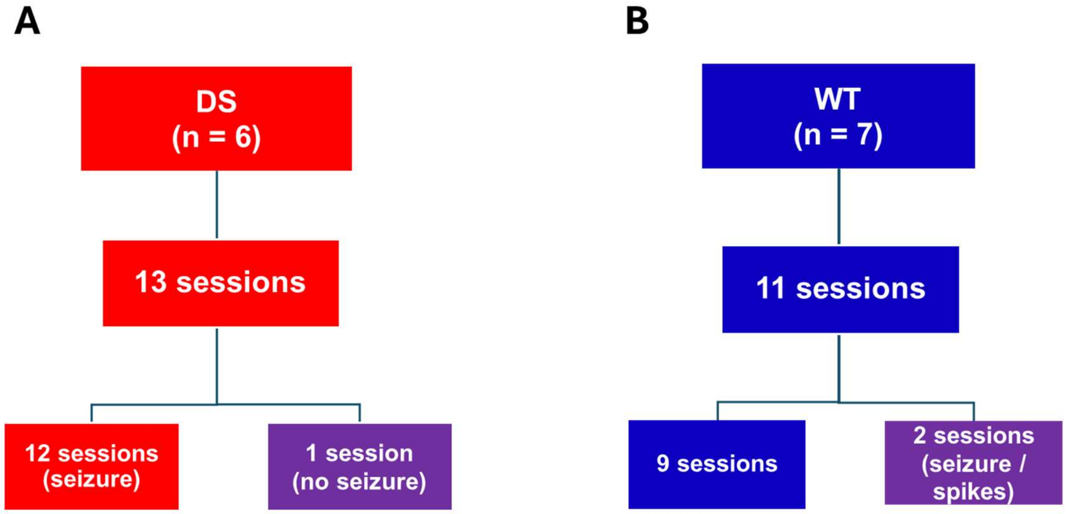


**(A)** Overview of fiber photometry data obtained from DS mice. All mice underwent at least two hyperthermia sessions. 1 session was analyzed separately as no seizure was observed.

**(B)** Overview of fiber photometry data obtained from WT control mice. Both sessions from 1 WT mouse were excluded from population analysis due to an unanticipated evoked seizure or sustained epileptiform discharges.

#### Figure S4. Unanticipated trials with failed seizure (DS) or evoked epileptiform activity (WT).

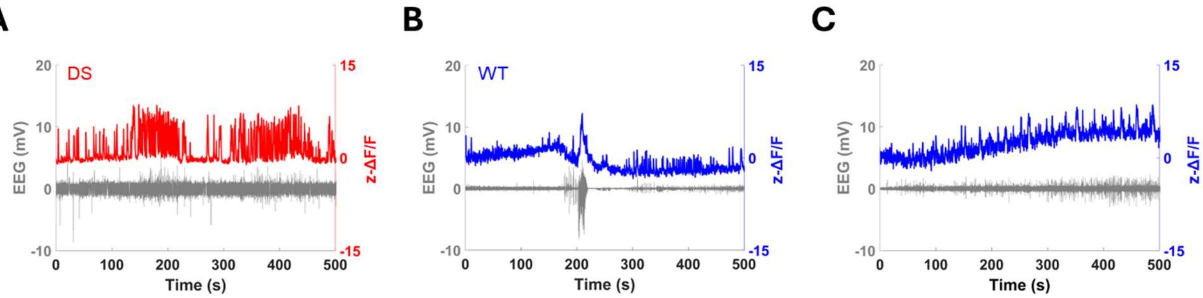


**(A)** EEG trace (mV; gray) overlaid with fiber photometry signal (z-ΔF/F; red) recorded from an DS mouse that did not have a seizure evoked by hyperthermia.

**(B)** EEG traces (mV; gray) overlaid with fiber photometry signals (z-ΔF/F; blue) recorded from a WT mouse that exhibited a seizure in the first hyperthermia session (left) and sustained spikes in the second (right).

#### Figure S5. Stereology sampling was adequate to achieve precise and accurate estimates of cell number.

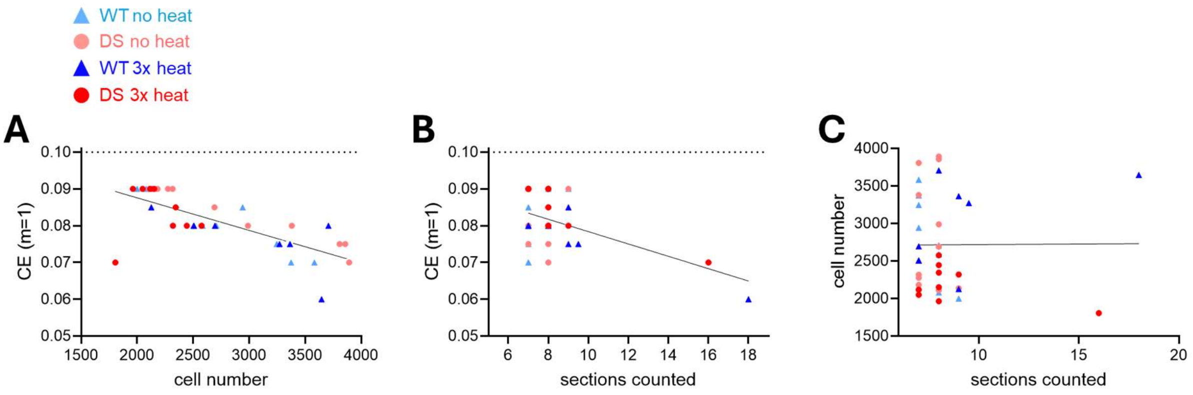


**(A)** Gundersen CE (m = 1) versus cPPN cell number estimate for each animal. A significant inverse relationship was observed ($R^2 = 0.54$, $F = 39.28$, $p < 0.0001$; simple linear regression), but all samples were below the coefficient of error cutoff of 0.1.

**(B)** Gundersen CE (m = 1) versus number of sections counted. A significant inverse relationship was observed ($R^2 = 0.25$, $F = 11.20$, $p = 0.002$; simple linear regression), but all samples were below the coefficient of error cutoff of 0.1. Note that for two brains, the Gundersen CE was > 0.1 when sampling every other section, so every section was sampled.

**(C)** There was no correlation between sections counted and cPPN cell number estimate for each animal ($R^2 = 3.026e\text{-}005$, $F = 0.0010$, $p = 0.97$; simple linear regression).

For all graphs, data points from WT mice were blue triangles and DS mice were red circles (open for no heat; closed for 3x heat).

#### Figure S6. Colocalization quantifications across independent experimenters.

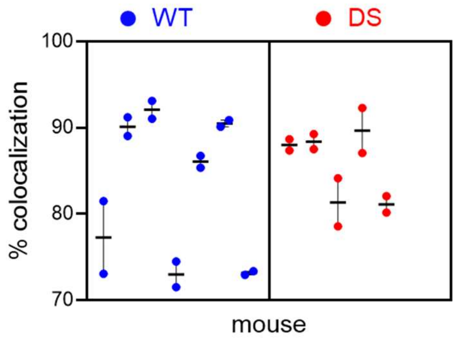


Colocalization was independently quantified by two investigators blinded to genotype for ChAT-tdT (n = 7) and DS-ChAT-tdT (n = 5) mice. Each column represents data from one mouse, with circles indicating the measurements from each experimenter. Nested t-test showed no significant difference (p = 0.55) between DS-ChAT-tdT (mean = 85.7) and ChAT-tdT (mean = 83.17) mice (p = 0.55). Data are presented as mean ± SEM.

### Tables

#### Table 1. cPPN stereology methodology.

| Condition | No heat | | 3x heat | |
|---|---|---|---|---|
| **Genotype** | **WT** | **DS** | **WT** | **DS** |
| **n (mice)** | 8 | 11 | 8 | 8 |
| **Section thickness (µm)** | 22.7 ± 0.7 | 24.0 ± 1.0 | 22.2 ± 1.9 | 19.4 ± 0.8 |
| **# of sections** | 7.6 ± 0.3 | 7.6 ± 0.2 | 9.3 ± 1.3 | 8.9 ± 1.0 |
| **% Difference between counts** | 8.3 ± 1.7 | 8.9 ± 1.2 | 7.9 ± 1.3 | 6.8 ± 1.7 |

Summary of stereological parameters in no heat and 3x heat groups, including the number of mice analyzed (n), section thickness, mean number of sections counted, and the percentage difference between two blinded independent counts. Data are presented as mean ± SEM.
